# Spatial correspondence in relative space regulates serial dependence

**DOI:** 10.1101/2023.06.27.546681

**Authors:** Jaeseob Lim, Sang-Hun Lee

**Affiliations:** Department of Brain and Cognitive Sciences, Seoul National University, 1 Gwanak-ro, Gwanak-gu, Seoul, Republic of Korea (08826)

## Abstract

Our perception is often attracted to what we have seen before, a phenomenon called ‘serial dependence.’ Serial dependence can help maintain a stable perception of the world, given the statistical regularity in the environment. If serial dependence serves this presumed utility, it should be pronounced when consecutive elements share the same identity when multiple elements spatially shift across successive views.

However, such preferential serial dependence between identity-matching elements in dynamic situations has never been empirically tested. Here, we hypothesized that serial dependence between consecutive elements is modulated more effectively by the spatial correspondence in relative space than by that in absolute space because spatial correspondence in relative coordinates can warrant identity matching invariantly to changes in absolute coordinates. To test this hypothesis, we developed a task where two targets change positions in unison between successive views. We found that serial dependence was substantially modulated by the correspondence in relative coordinates, but not by that in absolute coordinates. Moreover, such selective modulation by the correspondence in relative space was also observed even for the serial dependence defined by previous non-target elements. Our findings are consistent with the view that serial dependence subserves object-based perceptual stabilization over time in dynamic situations.

## Introduction

Objects in natural settings rarely alter their features. Even when alterations do occur, they are not random but exhibit statistical regularities over time^1,2^. For instance, sequentially appearing objects often have similar orientations^3^. Given this temporal steadiness of the natural environment, an ideal perceptual system would exploit this stability to accurately estimate object features. As an observation consistent with this view^2,4^, researchers consider a phenomenon called “serial dependence,” where our perceptual estimates tend to gravitate to the previous estimates^5,6^. Suppose the color perception of the current object is assimilated to the perceived color of the preceding object. Then, our perceptions of colors would become more stable than the colors actually are due to such serial dependence. This assimilation in feature perception across time appears to be a general phenomenon in the visual system, as it has been observed in other species^7^ and for various types of visual features. These features range from low-level features such as orientation, color, and spatial frequency to high-level features such as numerosity, ensemble statistics, facial expressions, and people identity^5,8–13^.

However, serial dependence does not always occur to the same degree, but its magnitude is influenced by many factors. One of the most fundamental and well-established factors is the spatial correspondence between consecutive elements: the current element is maximally attracted to the previous element that appeared in the same location and becomes progressively less attracted as the distance between them increases. This spatial specificity of serial dependence has been widely observed^5,14–17^ and even exploited as a basis for exploring other aspects of serial dependence^18–20^.

Having established spatial specificity as a fundamental property of serial dependence, one faces the critical issue of determining the coordinate system in which serial dependence is best characterized for its spatial specificity. Resolving this issue can provide important clues to understanding the utility and underlying mechanism of serial dependence. The specificity in world-centered coordinates would suggest that serial dependence helps stabilize our perception of ‘local features’ in the outer world—irrespective of where we look—and point to cortical areas representing features in world-centered space as its neural locus. The specificity in retinotopic coordinates would suggest that the utility of serial dependence is to stabilize our sense of an entire image formed in the retinae, and its neural locus resides in the early retinotopic visual areas. Using tasks where only a single element was presented or a single target was cued^5,14^, the spatial specificity of serial dependence was better characterized in retinotopic coordinates than in world-centered coordinates.

Here, as another coordinate system in which serial dependence is spatially specified, we considered the relative (object-centered) coordinate system^21–26^, which has not been investigated in the context of serial dependence. World-centered and retinotopic coordinates can both be considered ‘absolute’ coordinates in that the positions of elements are defined in reference to ‘temporally fixed points’ in any given space. By contrast, the positions of elements in relative coordinates are defined in reference to other elements within an object (e.g., “the nose is above the lips within a face”). It should be noted that the positions of elements remain unchanged in relative coordinates even when the reference items vary in absolute space over time.

In a static situation (Fig 1a), the spatial correspondence between any given two sequential items is indistinguishable in absolute or relative coordinates. However, their spatial correspondence becomes distinguishable in a dynamic situation where a group of elements (i.e., grouped objects or parts belonging to an object) change in position over consecutive views (Fig. 1b,c), as is often the case in natural settings. In the latter case (Fig. 1b,c), the sequential items matched in relative coordinates have the same identity, while those matched in absolute coordinates differ in identity. If the objective function of serial dependence is to stabilize our perception of the elements with the same identity (e.g., perception of objects or local parts within a single object) across time^27^, as previously claimed^9,15,28–30^, serial dependence should be routed in relative coordinates (Fig 1c) rather than in absolute coordinates (Fig. 1b) in a dynamic situation.

**Figure 1.**
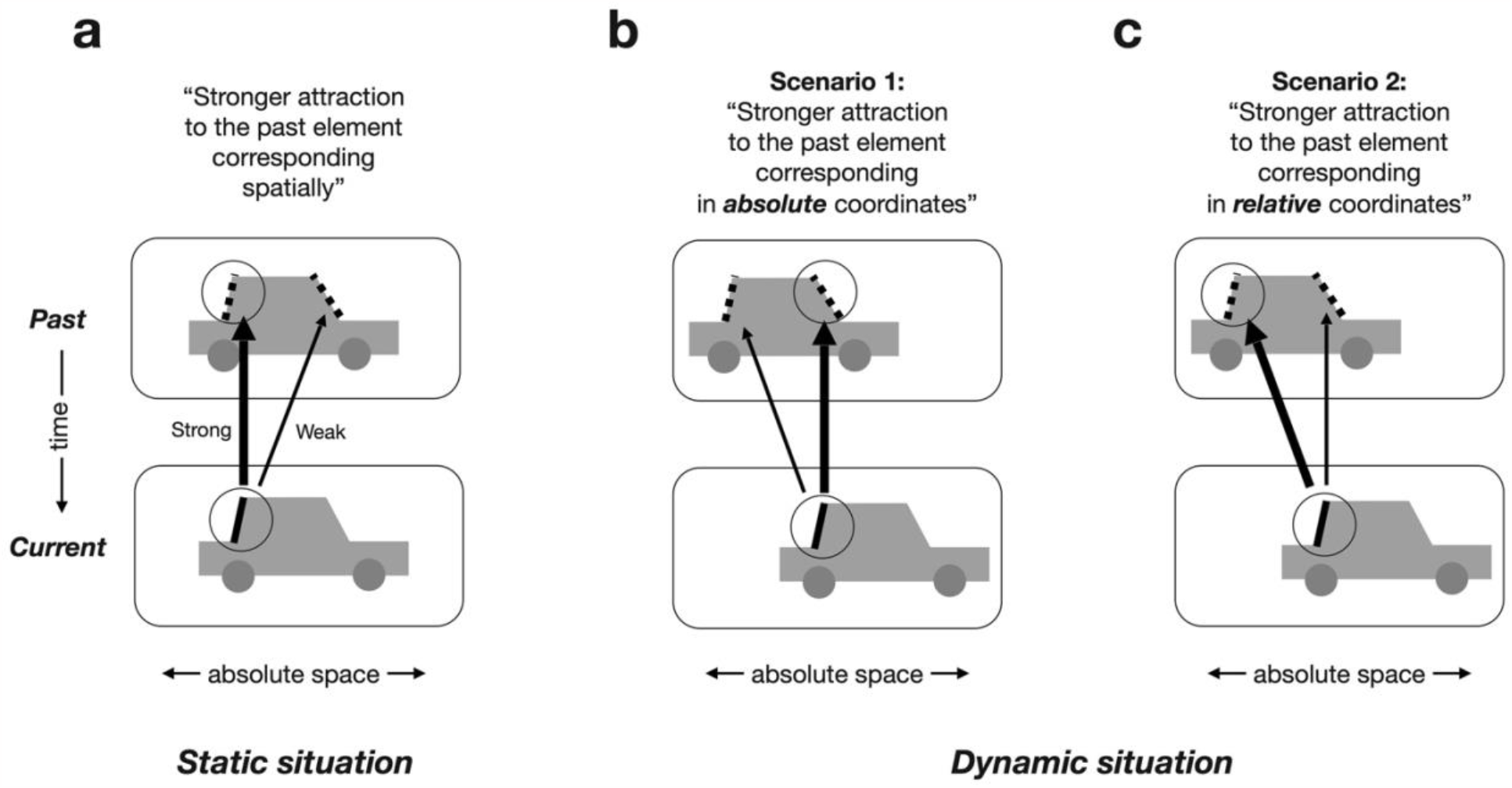
Spatial correspondence and attractive sequential effects in static (**a**) and dynamic (**b, c**) settings. Attractive effects (serial dependence) are known to occur preferentially between the consecutive elements with spatial correspondence (black circled elements in past and current). This means that attractive effects are stronger between spatially matched elements (thick arrow) than between unmatched elements (thin arrow). There are two types of spatial correspondence, one defined in absolute coordinates and the other in relative coordinates. These two types are indistinguishable when an object remains static across views (**a**). By contrast, the absolute and relative spatial correspondences can be distinguished when an object shifts in position across views (**b, c**). In such a dynamic setting, it is not the spatial correspondence in absolute coordinates (two circles in **b**) but spatial correspondence in relative coordinates (two circles in **c**) that determines whether consecutive elements match in identity. Thus, if serial dependence is transmitted between consecutive elements with the same identity, it is expected to be more pronounced between the elements corresponding in relative coordinates than those in absolute coordinates. The thickness of the arrows represents the magnitude of serial dependence.

Besides the rationale put forth above, prior empirical work where serial dependence was not directly investigated suggests that human performance in various perceptual tasks is influenced by the spatial correspondence defined in relative coordinates in similar dynamic situations. In a letter naming task, the priming effect in naming speed was determined by whether a target matched a primer in relative coordinates instead of absolute (retinotopic) coordinates^31^. In a visual search task, the pre-cueing effects, both in speed and accuracy, were also governed by the spatial proximity between the cue and target in relative coordinates^32^. In a task where motion perception requires temporal integration of multiple successive elements, the motion integration path conformed to the spatial proximity defined in relative coordinates^26,33^. Note that the tasks used in these studies, despite their differences in stimuli and responses, all entail the sequential effects in non-static situations. Because serial dependence can be considered a sequential effect, human reliance on the proximity defined in relative coordinates in the naming, search, and motion-perception tasks suggests that serial dependence is likely to transpire over the successive elements matched in relative coordinates when multiple elements undergo shifts as a group or as parts belong to the same object.

So far, we have provided the reasons to expect that the spatial specificity of serial dependence is better explained in relative coordinates than absolute coordinates. However, there are also good reasons to expect that the alternative scenario is at work. From a bottom-up, mechanistic perspective, the influence of preceding stimuli on ensuing stimuli may reflect the sensory-level carry-over effects induced by the lingering or reactivated traces of neural activity associated with the previous stimuli^34–36^. If such sensory-level mechanisms mediate serial dependence, then the spatial specificity of serial dependence will be better accounted for in retinotopic coordinates, as previously observed in a single-target situation^5,14^. Thus, it is an empirical matter whether serial dependence is regulated by the spatial correspondence in relative or absolute coordinates.

The lack of exploration into the spatial specificity of serial dependence in relative coordinates is a consequence of earlier research relying on task paradigms where only a single element is presented in each view^6,27^ or pre-cued even when presented with other elements^5,19,20^. Multiple ‘task-relevant’ elements—i.e., ‘targets’—are required to define the spatial correspondence between sequential items in relative coordinates. For that matter, a recent study^15^ reported the spatial specificity of serial dependence in a setup with two task-relevant targets displayed at each view. However, such specificity cannot be considered to signify the spatial specificity of serial dependence in relative coordinates since the two target locations remain fixed over successive views (like those shown in Fig. 1a). The spatial correspondence in relative and absolute coordinates can be distinguished only when multiple elements shift in location over successive views (Fig. 1b,c).

Here, we developed a task paradigm where two oriented grating elements shift their locations in unison over successive trials while maintaining their distance, and observers estimated the orientation of the retro-cued element on each trial (Fig. 2). This procedure is similar to the one used in the previous studies that demonstrated the effect of the proximity in relative coordinates on motion perception^26,33,37^. As they did, we promoted the grouping of two elements—so that the two elements are treated as two components belonging to a single object—by leveraging the ‘proximity between elements’ and ‘common fate’ rules of Gestalt^38–40^. With this task paradigm, we show that the orientation estimate of the current target is more strongly attracted to the orientation of the previous target matched in relative coordinates rather than the one matched in absolute coordinates. This suggests that human observers rely on spatial correspondence in relative coordinates in routing serial dependence between multiple elements over successive views to exploit environmental stability effectively.

**Figure 2.**
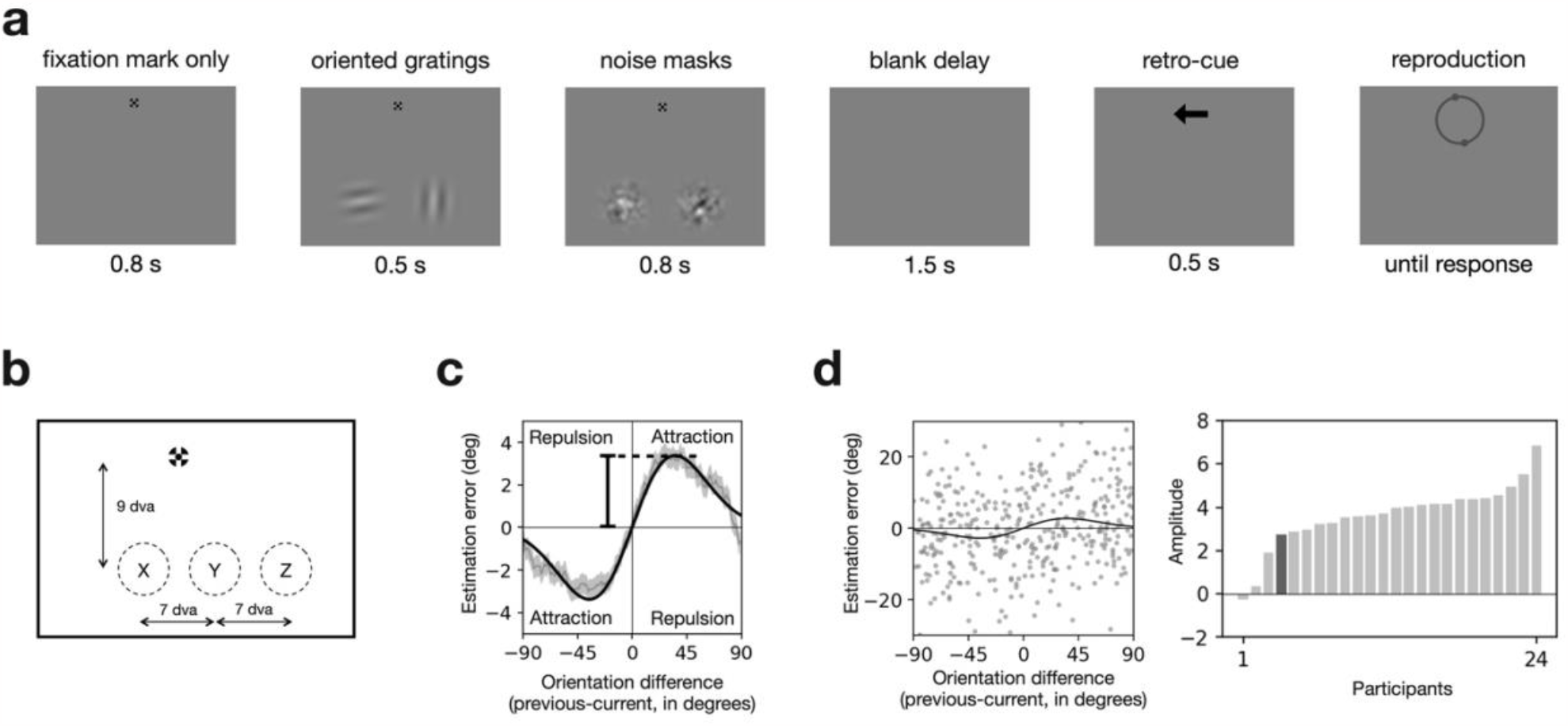
Experimental design and the definition of the strength of serial dependence. **a**, Trial structure. While fixating on a fixation mark, participants briefly viewed two Gabor patches in the lower visual field, which were followed by masks. After a short delay, they reported the orientation of the patch retro-cued by an arrow. **b**, Spatial arrangement of stimuli. On any given trial, two horizontally arranged Gabor patches appeared simultaneously. Gabor patches appeared randomly in two adjacent spots out of three spots (‘X and Y’ or ‘Y and Z’) beneath the fixation mark, which means the distance between the two patches was fixed across trials. **c**, Serial dependence curve merged across participants. The orientation estimation errors in the current trial were plotted against the orientation difference between the previous and current trials. The thin gray curves and shades represent the across-participant averages of the errors and their standard error of means (running average; window size = 20). The thick black curve is the Gaussian derivative fitted to the observed errors. The strength of serial dependence was quantified by the amplitude of the fitted Gaussian derivative (peak value; see Methods). **d**, Individual differences in serial dependence. The left panel plots the estimation errors of an example individual against the orientation difference between the previous and current trials. Gray dots represent trial-to-trial errors, and the black curve is the Gaussian derivative fitted to those errors. The right panel plots the strength of serial dependence for twenty-four individual participants in an ascending order. The dark bar represents the strength of serial dependence for the individual whose estimation errors are depicted in the left panel.

## Results

### Manipulating the spatial correspondence between successive targets in absolute and relative coordinates

Twenty-four human individuals participated in an experiment where they performed a delayed orientation estimation task on sequences of visual stimuli. On each trial, two oriented gratings simultaneously appeared and were then masked, and, after a delay, one of them was indicated by an arrow at the fixation mark (Fig. 2a). Participants were then required to reproduce the remembered orientation of the grating ‘retro-cued’ by that arrow. The retro-cued grating was chosen randomly on each trial. When reproducing the orientation of the retro-cued grating, participants were instructed to rotate a pair of dots on the circle around the fixation mark until the virtual line between the dots aligns with the remembered orientation of the target (Fig. 2a; see Methods for details). The two gratings were horizontally arranged beneath the fixation mark, and their centers were separated from each other by 7 degrees in visual angle (dva) (Fig. 2b). On each trial, the position of the paired gratings was randomly determined to be either the left two spots (marked as ‘X’ and ‘Y’ in Fig. 2b) or the right two spots (marked as ‘Y’ and ‘Z’ in Fig. 2b) of three retinotopically fixed spots. Throughout the experiment, participants kept fixating their gaze on a fixation mark whenever the mark was presented while their gaze position was monitored by a video-based eye-tracker. We excluded the trials from further analysis where a participant’s gaze deviated from the fixation mark by more than 1.5 dva during the fixation period (see Methods for details).

The above spatial arrangement and randomization of the gratings and retro-cue allowed us to separately manipulate the two variables of our interest: (i) the spatial correspondence between consecutive targets in absolute coordinates and (ii) that in relative coordinates. The consecutive targets (i.e., retro-cued gratings between consecutive trials) were apart by 0, 7, or 14 dva in absolute coordinates and by 0 or 7 dva in relative coordinates (Supplementary Fig. S1). To match the levels of spatial correspondence in absolute coordinates to those in relative coordinates, we excluded the trials from analysis where the consecutive targets were apart by 14 dva (marked as dashed boxes in Supplementary Fig. S1). As a result, the remaining trials could be sorted into two-by-two factorial conditions based on the spatial correspondence between consecutive targets in absolute and relative coordinates (Fig. 3a).

**Figure 3.**
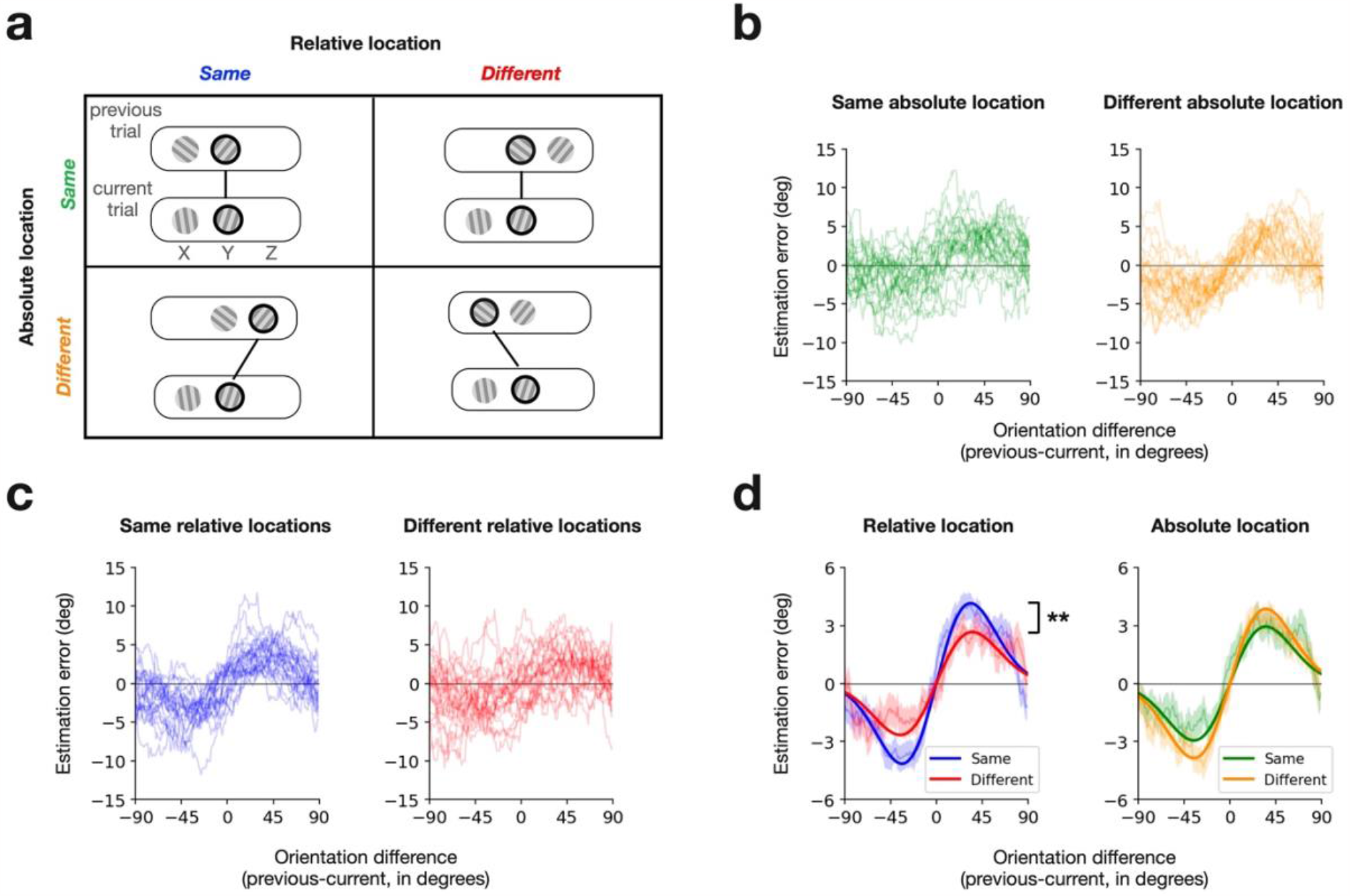
Effects of spatial correspondence on target-to-target serial dependence. **a**, Conditions of spatial correspondence between consecutive retro-cued targets in absolute and relative coordinates. Each panel shows an example sequence of displays in consecutive trials (top and bottom round rectangles for the previous and current trials, respectively), where the thick black lines and circles indicate the retro-cued targets. X, Y, and Z correspond to those shown in Fig. 2b, indicating the three possible spots for grating stimuli. In absolute coordinates, trials are sorted into the same-absolute-location (top row) or the different-absolute-location (bottom row) conditions. In relative coordinates, trials are sorted into the same-relative-location (left column) or the different-relative-location (right column) conditions. **b**,**c**, Serial dependence curves from individual participants for the spatial correspondence conditions defined in absolute (**b**) and relative (**c**) coordinates. Each panel plots the running averages (window size of 20°) of the estimation errors in the current trial as a function of the difference in target orientation between the previous and current trials. Thin curves represent individual participants, and their colors correspond to the correspondence conditions defined in **a. d**, Comparison of the strength of serial dependence between the spatial correspondence conditions defined in absolute (right) and relative (right) coordinates. Each panel shows the serial dependence curves merged across all participants for the two correspondence conditions. The thin curves and shades represent the across-participant averages of the errors and their standard error of means. The thick curves represent the Gaussian derivatives fitted to the observed errors, quantifying the magnitude of serial dependence with amplitudes. The two stars indicate that serial dependence significantly (p<0.01) differed in amplitude between the same-relative-location and different-relative-location conditions.

### Serial dependence in the data merged across conditions

We began our analysis by assessing whether participants’ delayed orientation estimates follow the previously reported pattern of serial dependence. We merged the data from all the participants across the correspondence conditions and plotted their estimation errors on the current target as a function of its orientation difference from the previous target. The running averages of the errors (window size of 20°) followed a clockwise-rotated ‘S’ shape (gray curve with a shade in Fig. 2c), which is the canonical pattern of serial dependence. Following the convention, we fitted the Gaussian derivative to the errors (thick black curve in Fig. 2c; Pearson correlation r = 0.985) and quantified the strength of serial dependence by the amplitude of the fitted Gaussian derivative (vertical bar in Fig. 2c; 3.38° with bootstrap-based 95% CI of [2.96°, 3.85°]; see Methods for details). When the same amplitude estimation procedure was applied to the data at an individual level (Fig. 2d, left panel), all participants except for one showed positive values of amplitude (Fig. 2d, right panel; range in degree, [-0.27°, 6.86°]; mean, 3.63°; SD, 1.45°). This indicates that the orientation estimate of the current target tended to be attracted to the orientation of the previous target for the vast majority of participants.

Having confirmed the presence and prevalence of serial dependence in the merged data, we turned to the data conditioned on the spatial correspondence in absolute and relative coordinates. They were compared in the strength of serial dependence based on the same procedure used in the merged data, which has been established as a reliable way to compare the amplitudes between two experimental conditions^12,15,19,27^.

### Effects of spatial correspondence on target-to-target serial dependence

To assess how the serial dependence of the current target on the previous target is influenced by their spatial correspondence in absolute coordinates, we split and merged the trials into the same-absolute-location and different-absolute-location conditions (top versus bottom rows in Fig. 3a) and plotted the errors against the orientation difference between the consecutive targets (Fig. 3b). Likewise, the same trials were split and merged into the same-relative-location and different-relative-location conditions (left versus right columns in Fig. 3a; Fig. 3c) to assess the impact of the spatial correspondence in relative coordinates on serial dependence.

The spatial correspondence significantly affected the strength of serial dependence in relative coordinates but not in absolute coordinates (Fig. 3d). In relative coordinates, the estimated amplitude of serial dependence was substantially greater in the same-relative-location condition (4.14°; bootstrapbased 95% CI, [3.48°, 4.89°]) than in the different-relative-location condition (2.66°; bootstrap-based 95% CI, [1.93°, 3.42°]) (p = 0.004, permutation test). By contrast, when measured in absolute coordinates, it was slightly smaller in the same-absolute-location condition (2.94°; bootstrap-based 95% CI, [2.27°, 3.68°]) than in the different-absolute-location condition (3.85°; bootstrap-based 95% CI, [3.26°, 4.57°]), although the difference between the two conditions was not statistically significant (p = 0.07, permutation test).

It should be noted that the two correspondence conditions defined in relative coordinates were not perfectly matched in the ratio of the same-absolute-location trials to the different-absolute-location trials. Specifically, the same-absolute-location and different-absolute-location trials were balanced in number for the same-relative-location condition but not for the different-relative-location condition (Supplementary Fig. S1). Could this relatively smaller proportion of the same-absolute-location trials in the differentrelative-location condition have contributed to the spatial correspondence’s observed effect on serial dependence in relative coordinates? For this to be true, the strength of serial dependence in the sameabsolute-location condition should be greater than that in the different-absolute-location condition. However, as summarized above (Fig. 3b,d), the strength of serial dependence was actually weaker— insignificantly though—in the same-absolute-location condition than in the different-absolute-location condition. Thus, the effect of spatial correspondence in relative coordinates on serial dependence cannot be attributed to the different ratio of the same-absolute-location trials to the different-absolute-location trials between the two correspondence conditions in relative coordinates.

### Effects of spatial correspondence on non-target-to-target serial dependence

Our retro-cueing procedure (Fig. 2a) requires participants to retain orientation information for both gratings during the delay period because a target grating cannot be identified until indicated by the retro-cue. As a result, the orientation of the non-target stimulus in the previous trial was not overtly reported, but it might still influence the perception or memory of the target orientation in the current trial. Therefore, serial dependence might also occur between the previous non-target and the current target. We indeed confirmed that the orientation estimates of the current target were attracted to the orientation of the previous nontarget when all data were merged across participants (Fig. 4a; 0.87° with bootstrap-based 95% CI of [0.40°, 1.81°]). We will call this the ‘non-target-to-target’ serial dependence to distinguish it from the ‘target-to-target’ serial dependence described in the previous section. The presence of the non-target-to-target serial dependence prompts a question of whether its strength would still be influenced by the spatial correspondence in relative coordinates.

**Figure 4.**
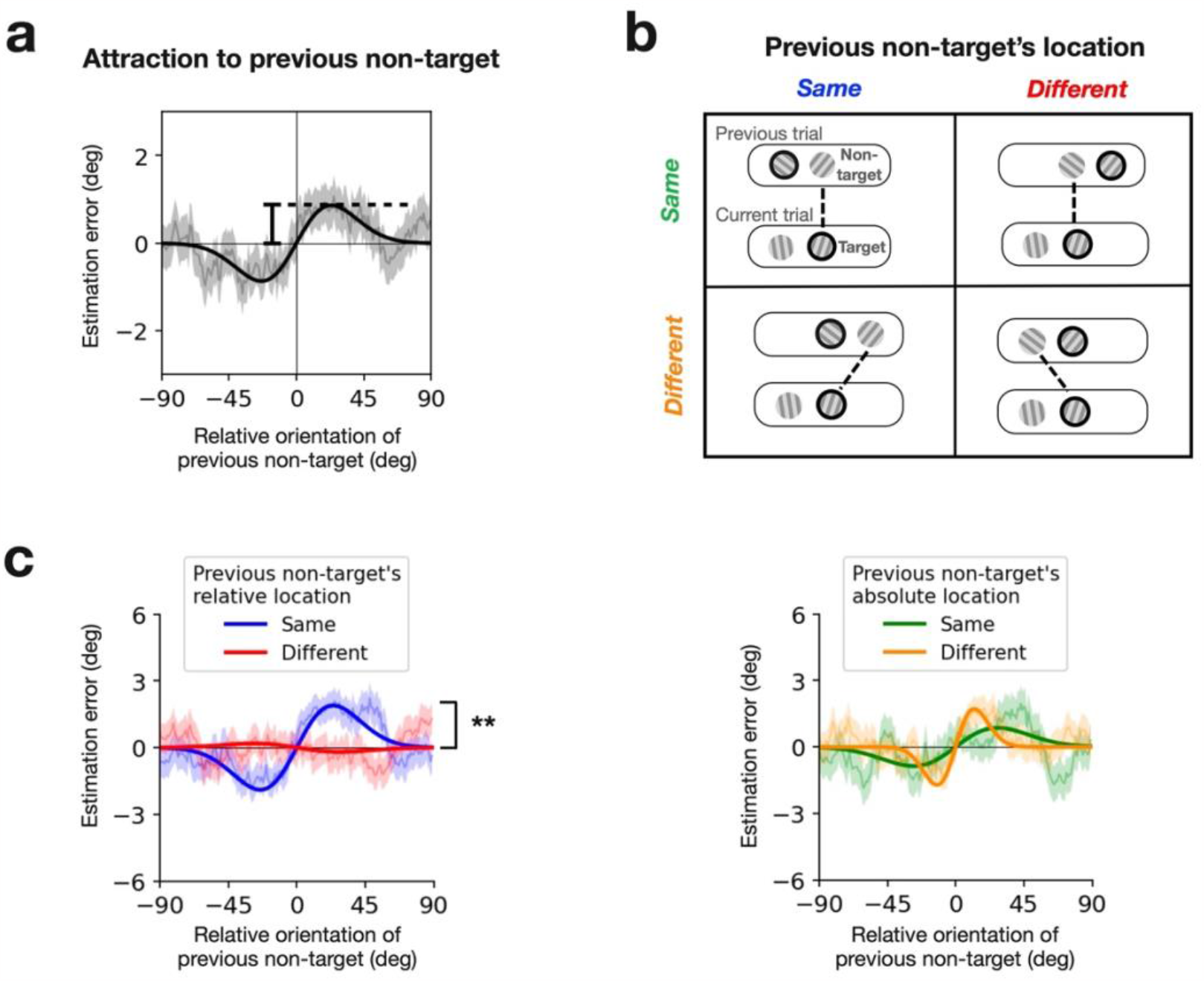
Effects of spatial correspondence on non-target-to-target serial dependence. **a**, Serial dependence curve merged across participants. The orientation estimation errors in the current trial were plotted against the orientation difference between the non-target in the previous trial and the target in the current trial. The thin gray curves and shades represent the across-participant averages of the errors and their standard error of means (running average; window size = 20). The thick black curve is the Gaussian derivative fitted to the observed errors. The strength of serial dependence was quantified by the amplitude of the fitted Gaussian derivative (as indicated by the vertical bar). **b**, Conditions of spatial correspondence between the non-target in the previous trial and the target in the current trial in absolute and relative coordinates. Each panel shows an example sequence of displays in consecutive trials (top and bottom round rectangles for the previous and current trials, respectively), where the thick black lines and circles indicate the non-target in the previous trial and the target in the current trial. Otherwise, the format is identical to that of Fig. 3a. **c**, Comparison of the strength of serial dependence between the spatial correspondence conditions defined in absolute (right) and relative (right) coordinates. The format and symbols are identical to those of Fig. 3a.

To answer this question, by applying the same procedure used for the target-to-target serial dependence (Fig. 3a), we split and merged the trials into the two spatial correspondence conditions, either in absolute coordinates (top versus bottom rows in Fig. 4b) or in relative coordinates (left versus right columns in Fig. 4b). Also, as was done for the target-to-target serial dependence, we did not include in analysis the trials where the current target and non-target were apart by 14 dva in absolute coordinates.

Results were qualitatively similar to what we observed in the target-to-target serial dependence: the spatial correspondence significantly affected the strength of serial dependence in relative coordinates but not in absolute coordinates (Fig. 4c). In relative coordinates, the estimated amplitude of non-target-to-target serial dependence was substantially greater in the same-relative-location condition (1.89°; bootstrap-based 95% CI, [1.20°, 2.87°]) than in the different-relative-location condition (-0.19°; bootstrap-based 95% CI, [- 1.72°, 1.23°]) (p = 0.002, permutation test). By contrast, when measured in absolute coordinates, there was no significant difference in the strength of the non-target-to-target serial dependence between the sameabsolute-location condition (0.86°; bootstrap-based 95% CI, [0.04°, 2.00°]) than in the different-absolute-location condition (1.714°; bootstrap-based 95% CI, [0.69°, 3.14°]) (p = 0.286, permutation test). These results suggest that serial dependence selectively occurs between the successive elements matched in relative coordinates as long as the previous element’s feature is retained in memory, irrespective of whether it was overtly reported or not.

## Discussion

Guided by the view that serial dependence reflects the perceptual system’s strategy of leveraging environmental stability to estimate object features, we hypothesized that serial dependence preferentially occurs between successive elements matched in relative coordinates when multiple elements shift in position over time. Consistent with this hypothesis, we found that the spatial correspondence in relative coordinates increased the strength of serial dependence. By contrast, such an increase was not found when the spatial correspondence was defined in absolute coordinates.

There are two different perspectives to consider when explaining how the spatial location of elements impacts serial dependence depending on whether it relates to serial dependence based on ‘local features’ or ‘element identities.’ The ‘feature-based’ view focuses on the statistical regularity governing the spatiotemporal relationship among local features in the environment. In this view, as consecutive elements get closer in space, they tend to have similar features, which leads to an increase in serial dependence. Here, the spatial proximity between consecutive elements directly regulates serial dependence, irrespective of their identity being the same or different. The feature-based view is well captured by the concept of the “continuity field”^5^, which assumes that the perceptual system strives to achieve visual perception with spatiotemporal stability in terms of local features, where element identity does not play any role. By contrast, in the ‘identity-based’ (or ‘object-based’) view, the primary reason for the serial dependence pronounced by spatial correspondence is that spatial correspondence between consecutive elements is an indicator that those elements share the same identity^41,42^. So, in this view, in a situation where spatial correspondence cannot play a role as an identity indicator, serial dependence will not increase by spatial correspondence. In the current work, by distinguishing between two kinds of spatial correspondence, one irrelevant to identity but more informative about local features (correspondence in absolute coordinates) and the other informative about item identity (correspondence in relative coordinates), we could demonstrate that the latter type of spatial information regulates serial dependence when multiple potential targets shift in position over consecutive views. Therefore, our findings support the identity (object)-based view of serial dependence, where the spatial correspondence between successive items informs the system that they share the same identity, and thus, their features are unlikely to change abruptly.

We found that the magnitude of serial dependence substantially dropped as the distance between consecutive targets in relative coordinates only increased from 0 dva to 7 dva (from 4.14° to 2.66°, reduction of 1.48° for the target-to-target serial dependence; from 1.89° to -0.19°, reduction of 2.08° for the non-target-to-target serial dependence). Such a marked reduction due to a slight mismatch in space is hard to explain based on the effects of spatial proximity on serial dependence in previous studies. In those studies, the magnitude of serial dependence did not decrease substantially until the consecutive targets became apart by 10 dva in absolute coordinates^5,14,19^. According to our view, which is that the spatial correspondence in relative space functions as a pointer to route assimilation “selectively” between the consecutive elements sharing the same identity among multiple potential targets, our task paradigm requires high spatial specificity due to the proximity of multiple targets. By contrast, in the task paradigms of previous studies, a single element or a single attended (pre-cued) target—exists at each view, and its identity is not required to be spatially separated from other co-existing items. We thus conjecture that the spatial specificity of serial dependence is not fixed but is likely to vary substantially depending on the functional role of spatial location in a given task paradigm. The spatial specificity is likely to be pronounced when it serves as a guide to selectively route assimilation between the elements belonging to a single object across time.

As stated earlier on, the current work was primarily motivated by the ‘object-based’ view on serial dependence, which predicts that serial dependence would be preferentially routed between the consecutive elements spatially matched within a single object. Guided by this view, to make the two neighboring elements appear to belong to a single object, we purposefully exploited the ‘proximity’ and ‘common-fate’ rule of Gestalt^38–40^ by shifting them simultaneously while keeping them at a fixed short distance (7 dva) throughout the experiment. This manipulation mimics a natural setting where a single object consisting of multiple elements stay (reappearing in the same spot in absolute space, as depicted in Fig. 1a) or jump (appearing in a different spot in absolute space, as depicted in Fig. 1b,c). Under that setting, we confirmed the prediction of the object-based view of serial dependence. Having interpreted our findings as such, we conjecture that the effect of the correspondence in relative coordinates on serial dependence is likely to be substantially weakened or disappear in a setting where two elements are no longer perceived as a group or parts belonging to a single object, such as when they were far apart, or their distance varied across trials^38–40^. Further investigation is required to identify the situations and factors facilitating and suppressing the spatial specificity of serial dependence in relative coordinates.

When spatial correspondence was defined in absolute coordinates, serial dependence was expected to be greater when two consecutive elements were matched in absolute coordinates than when they were not, based on its spatial specificity in absolute coordinates reported in previous studies^5,14,19^. However, the strength of serial dependence in the same-absolute-location condition did not significantly differ from that in the different-absolute-location condition (the right panels in Fig. 3d and Fig. 4c). We conjecture that in the same-absolute-location condition, the attraction of the current orientation estimation to the previous stimulus orientation could have been attenuated by a repulsive sequential effect^43^, which selectively occur between stimuli matched in absolute coordinates. Our conjecture aligns with a recent body of work supporting the view that the ‘attraction’ and ‘repulsion’ types of sequential effects have distinct origins yet coexist^19,20,44–48^.

Our findings should be distinguished from the debates about whether the spatial tuning of serial dependence is governed by the distance in retinotopic coordinates^14^ or in world-centered coordinates^5^. In our terms, both types of distance are considered the distance in absolute coordinates. Instead, our findings suggest that serial dependence is regulated by the distance between consecutive targets in object-centered coordinates in dynamic settings with multiple targets. The object-centered space is similar to the world-centered space in that both use a reference frame outside the observer, unlike the ego-centric retinotopic space. However, the world-centered space’s reference frame is a fixed point that remains constant over time in the world, whereas the object-centered space’s reference frame is part of an object^21–24^. Given that the brain initially processes spatial information in the ego-centric retinotopic space, representing spatial information in the object-centered space requires extra calculations to relocate elements within a new reference frame. Such extra calculations are known to occur in high-level associative cortical regions, including the supplemental eye field^25^, parietal cortex^22^, and lateral occipital-temporal cortex^21^. Our work indicates that the serial dependence between successive local features is controlled in the object-centered space, suggesting that the neural mechanism responsible for serial dependence is located in those associative cortices, or at least entails the translation of spatial information from retinotopic to object-centered coordinates. This possibility seems consistent with the view that serial dependence is more likely to emerge at stages higher than the retinotopic stage where adaptation-based repulsive sequential effects are dominant ^45,48,49^.

Our study highlights the utility of spatial correspondence in relative coordinates as a factor modulating the strength of serial dependence in dynamic situations. To be sure, our task paradigm is obviously limited in covering all possible dynamic situations where serial dependence is regulated by spatial correspondence in relative coordinates. For example, we created a dynamic situation by shifting elements across views while asking participants to fixate their gaze on a fixed spot in absolute space. Alternatively, another dynamic but retinotopically identical situation can be created by asking them to shift their gaze while keeping the elements in the same place in absolute space. It is unclear whether spatial correspondence in relative coordinates affects serial dependence in this scenario, especially because it has been previously shown that retinal motions induced by saccades and those by real movements are processed differently^50^, and saccades themselves influence visual perception^51^.

In summary, we demonstrated that the correspondence between consecutive elements in the object-centered space augments their serial dependence. This supports the idea that serial dependence reflects the perceptual system’s strategy to stabilize its perception of sequential elements that share the same identity in natural settings where objects consisting of multiple components stay or move around.

## Methods

### Participants

Participants with normal or corrected-to-normal vision were recruited through offline and online postings on the Seoul National University internet community. Individuals unsuitable for eye tracking (e.g., wearing thick glasses) were excluded from the study. A total of twenty-four individuals (12 females), between the ages of 19-33 years (average age of 25.5), participated in the study. Each participant participated in a 90-minute experiment, received a compensation of 25,000 KRW, and gave written informed consent before the experiment. The experiment was conducted in compliance with the safety guidelines for human experimental research, as approved by the Institutional Review Board of Seoul National University (IRB No. 2108/001-012).

### Stimuli

Visual stimuli were Gaussian-enveloped sinusoidal gratings (Gabor patches) with a Michelson contrast of 0.3. To promote effective fixation, a bullseye with a cross-hair inside was used as a fixation marker^52^. We used a 24-inch LCD monitor (LG 24MP58VQ) to display the stimuli. The Gabor patches had a grating with a spatial frequency of 0.588 cycles per dva and a Gaussian envelope with a standard deviation of 1.1 dva. The centers of the two Gabor patches were fixed at a horizontal distance of 7 dva. Both patches were positioned 9 dva below the fixation marker. On each trial, the midpoint of the two patches was randomly placed either 0 dva or 7 dva to the right of the fixation marker (as shown in Fig. 2b). To make noise masks, we first low-pass filtered a pixel-by-pixel white noise pattern using a 2D Gaussian filter (a standard deviation of 0.2 dva) and applied a spatial 2D Gaussian filter (a standard deviation of 1.1 dva) to the low-pass filtered noise pattern. The contrast of the noise masks was normalized to a Michelson contrast of 0.9. The font size of the retro-cue arrow was 1 dva, measured in pixels. To report the orientation of the target, observers rotated two gray dots (each with a 0.5 dva) along a thick circle with a 2.5 dva around the fixation marker. The fixation marker, retro-cue, and orientation-reporting dots appeared in fixed positions on the upper center region of the screen (Fig. 2a). On each trial, the orientations and phases of the two gratings were randomly selected from the angles ranging from 0° to 179°, with a 1° increment. The stimuli were generated using PsychToolbox for MATLAB^53^.

### Task

Each participant performed 9 blocks of trials, with each block containing 45 trials. Each trial began with the fixation marker appearing on the upper center region of the screen. After a delay of 800 ms, two gratings appeared for 500 ms, followed by noise patches that lasted 800 ms. Participants were required to remember the orientations of both gratings for a delay of 1500 ms before the retro-cue indicated one of the two gratings as a target. The retro-cue was an upward or downward arrow that appeared for 500 ms and indicated the location of the target whose orientation was to be reported. Once the retro-cue disappeared, participants reported the orientation of the retro-cued target grating by rotating the reporting dots around the fixation marker. They used two keys on a keyboard to rotate the reporting dots and had to complete their report within a 10-s time limit. Once participants confirmed their report, the reporting dots vanished, and the fixation marker for the next trial appeared after a delay of 500 ms.

Participants’ gaze was monitored during stimulus presentation with a video-based eye-tracker.When their gaze significantly deviated from the fixation marker (see the “Eye movement tracking” below for details), a message reading “Your eyes have moved” was displayed below the fixation marker for 1 s to alert participants. When they failed to maintain their gaze fixation for three trials in a row, we prompted them to pay more attention and re-calibrated the eye-tracker.

### Eye movement tracking

Eye movement tracking was conducted using a desktop-mounted Eyelink 1000 system (SR research, Ottawa, Canada) at a sampling rate of 1000 Hz. To ensure accuracy, the eye-tracker was calibrated using a built-in 5-point routine (HV5) at the beginning of each block of trials, If participants’ fixation failed for three consecutive trials, the eye-tracker was re-calibrated. A fixation failure was defined by the following procedure. The average of the gaze position samples during the 240 ms before stimulus onset was calculated to determine the baseline gaze position for each trial. If any gaze position sample deviated by more than 1.5 dva from this baseline, the trial was considered a fixation failure. We confirmed that the gaze deviations were quite small and did not vary depending on where the gratings were located (Supplementary Fig. S2). Additionally, trials where eye blinks occurred during the fixation or stimulus presentation periods were also classified as fixation failure.

### Discarding trials

Outlier trials with large estimation errors were discarded from the analysis. To identify these outliers, we used Tukey’s method at an individual level by excluding trials in which estimation errors were below the lower quartile (Q1) minus 1.5 times the interquartile range (IQR) or above the upper quartile (Q3) plus 1.5 times the IQR. As a result, 5.15% of total trials were discarded. As previously stated, we excluded the fixation failure trials, which accounted for 2.75% of the total trials. We also removed trials with a target distance of 14 dva between consecutive targets in absolute coordinates, which accounted for 12.25% of the total trials. These trials were not included in the main data analysis.

### Data pooling across participants

Compared to other studies where only one target was remembered, orientation estimates tended to be noisier in our study, where two targets had to be remembered. This could be due to an increase in memory load^54^, contextual interferences from nearby stimuli^43,55^, or pronounced stimulus-specific biases^56^. As the estimation noise was high, it was not reliable to fit the serial dependence curve to individual data for each condition. Therefore, we pooled the data across individuals after demeaning the errors within individuals for statistical tests and visualization of serial dependence, following previous studies^12,15,19,27^.

### Curve fitting

We fitted the Gaussian derivative, 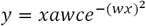, to the pooled data for each condition, where *y* is the estimation error; *x* is the previous target’s orientation relative to the current target’s orientation; *c* is a constant 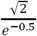; *a* and *w* are the free parameters associated with curve amplitude and width, respectively^5^. The best-fit parameters were determined by finding the values *a* and *w* that minimize the sum of squared errors with the Broyden, Fletcher, Goldfarb, and Shanno (BFGS) algorithm. To avoid overfitting and promote fast saturation, we bounded the values of *a* and *w* within [-50, 50] and [0.02, 0.2], respectively. The best-fit value of *a* was used as the quantity representing how much the current target’s orientation is attracted to the previous target’s orientation. We estimated the reliability of the best-fit parameter value by defining 95% confidence intervals using the following non-parametric bootstrap procedure. First, the trials in a given condition were randomly resampled with replacements from each individual’s data. Second, the resampled data were pooled across individuals. Third, the best-fit value of *a* was estimated by fitting the Gaussian derivative to the pooled data for each condition. We repeated this procedure 5,000 times to obtain the distribution of best-fit values and defined the 95% confidence interval based on this distribution.

### Statistical test

To evaluate the significance of the difference in serial dependence amplitude between any given two conditions, we carried out a permutation test^19,27^ as follows. First, the labels of the two conditions (e.g., same-relative-location and different-relative-location conditions) were shuffled for each individual data. Then, the individual data were pooled across participants according to the shuffled condition labels. Next, the best-fit value of *a* was estimated by fitting the Gaussian derivative function to the pooled data for each of the two shuffled-label conditions. Lastly, we calculated the difference in the best-fit value of *a* between the two conditions. We repeated this procedure 10,000 times to obtain the distribution of these ‘permutated’ differences and calculated the p-value of the observed (non-shuffled) difference in serial dependence amplitude based on this distribution.

## Acknowledgements

This research was supported by the Research Grant from Seoul National University (339-20220013), Brain Research Program Grants NRF-2021R1F1A1052020 and NRF-2017M3C7A1047860, and Basic Research Laboratory Program Grant NRF-2018R1A4A1025891 through the National Research Foundation of Korea (NRF) funded by the Ministry of Science and Information and Communications Technology.

## Author contributions

J.L. and S.L. designed the experiments, analyzed results, and wrote the manuscript. J.L. collected data.

## Competing Interests Statement

The authors declare no competing interests.

## Notes

### Competing Interest Statement

The authors have declared no competing interest.

### Summary of Updates

Previous experiment 1 and related parts were excluded from the manuscript. All figures were updated, too. Introduction and discussion is also revised.

