## Supplementary Fig. for "Spatial correspondence in relative space regulates serial dependence"

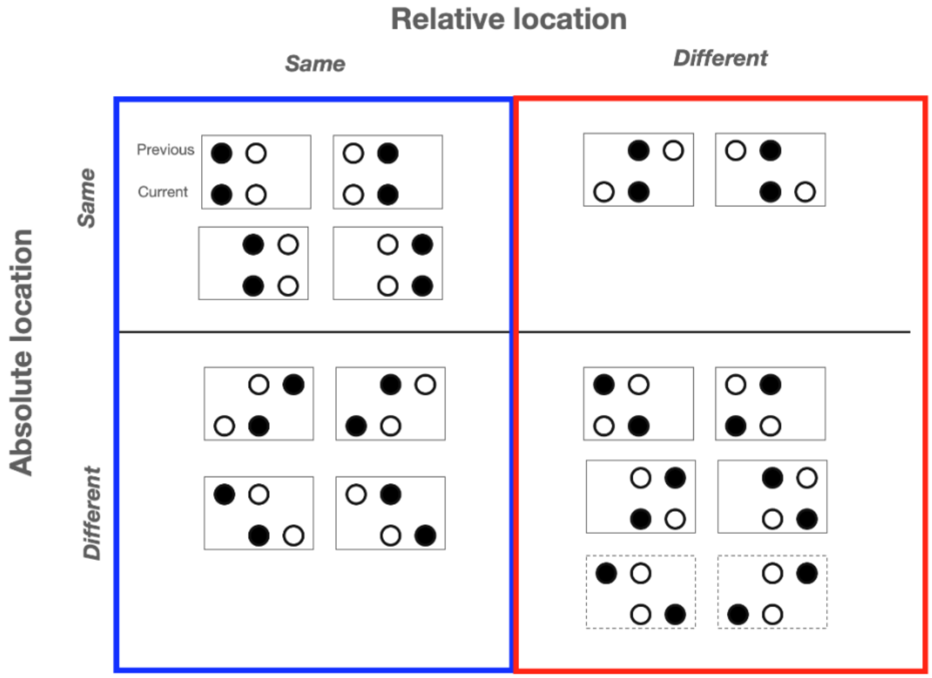


**Figure S1. Possible stimulus locations for the spatial correspondence conditions.** Each cell of the 2 x 2 table shows all possible combinations of stimulus locations for a given spatial correspondence condition. Within each box outlined by a thin line, filled and open circles demarcate the target and non-target locations on the previous (top) and current (bottom) trials. Because target and non-target locations were completely randomized, all sixteen possible combinations of target and non-target locations were equally probable (1/16). However, we excluded the trials where the consecutive targets were apart by 14 dva (those demarcated by the dashed boxes in the bottom right cell) from analysis to match the levels of target-to-target distance between the absolute-coordinate and relative-coordinate conditions

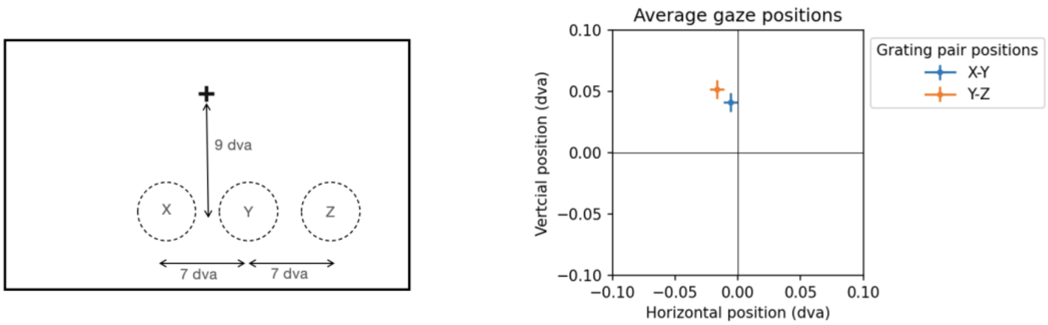


**Figure S2. Average gaze positions during stimulus presentation.** The left panel shows the three spots where grating stimuli appeared in the experiment. In any given trial, the two adjacent gratings appeared either in the X and Y locations or in the Y and Z locations. The right panel displays how much the gaze positions during stimulus presentation deviated from those during the 250 ms before stimulus onset. The blue and orange dots indicate the average deviations in gaze position when the gratings appeared in the X and Y locations and in the Y and Z locations, respectively. The horizontal and vertical bars correspond to the standard errors of the means. In our analysis, we excluded trials from analysis where gaze deviation was greater than 1.5 dva. To check whether gaze deviation varied depending on where the gratings were located, we compared the gaze deviation between the two location conditions (‘X and Y’ versus ‘Y and Z’). We found that the gaze deviations were quite small and did not differ when the grating appeared in the X and Y locations (-0.006 (mean) ± 0.006 (s.e.m.) dva for horizontal gaze deviation; -0.041 ± 0.008 dva for vertical gaze deviation) and when the grating appeared in the Y and Z locations (-0.017 ± 0.005 dva for horizontal gaze deviation; -0.052 ± 0.008 dva for vertical gaze deviation) ($t = 1.418$, $p > 0.1$ for horizontal gaze deviation; $t = 0.996$, $p > 0.1$ for vertical gaze deviation).
